# Subtractive and Divisive Methods for Isolating Neural Oscillations From Broadband Activity Yield Different, Even Conflicting, Power Estimates

**DOI:** 10.64898/2026.11.14.766643

**Authors:** Máté Gyurkovics, Emily Cunningham, Kathy A. Low, Grace M. Clements, Patrycja Kałamała, Diane M. Beck, Bradley Voytek, Monica Fabiani, Thomas Donoghue, Gabriele Gratton

**Author notes:** These authors contributed equally to this manuscript as first authors (Máté Gyurkovics, Emily Cunningham).

## Abstract

Understanding the role of oscillatory brain activity in cognition is a key objective in cognitive neuroscience. However, quantifying the power of narrowband neural oscillations is made challenging by the presence of non-oscillatory broadband activity in the signal. Recently, there has been a proliferation of methods designed to separate oscillatory from non-oscillatory (aperiodic) signals. To perform this separation, these methods must model the relationship between oscillatory and aperiodic components. Crucially, however, such methods rarely explicitly clarify whether they assume that this relationship is additive or multiplicative, an issue compounded by the common use of log-transformed data (i.e., subtraction in log space is equivalent to division in linear space). Here, we show the importance of justifying which assumption about component relations is applied. Using simulated power spectra with additive oscillatory and aperiodic components, we illustrate that methods that assume a multiplicative model, such as the popular *specparam* (formerly *fooof*) toolbox, systematically mis-estimate oscillatory activity in ways that can obscure real oscillatory differences between conditions, or even reverse their sign, creating an illusory effect. We then demonstrate these effects in two complementary empirical datasets: (a) a cross-sectional, between-subjects analysis of age-related differences in alpha power, showing how separation approaches impact effect size and detectability; and (b) a within-subjects analysis of pre-stimulus changes in alpha power in a detection task, showing how switching between approaches can greatly reduce the evidence for an effect, leading to different conclusions. These results highlight that researchers must explicitly report and justify assumptions about aperiodic and oscillatory signal combination, as these assumptions have implications not only for analyses of oscillatory activity and resultant conclusions, but also for integration of results across reports (which may implicitly employ different assumptions), and ultimately for the development of theories regarding electrophysiological features and their putative relations to cognition.

## 1. Introduction

Understanding the role of neural oscillations in cognition has long been a central aim of cognitive neuroscience (e.g., Buzsaki & Draguhn, 2004). In recent years, there has been a considerable increase in interest in separating oscillatory from broadband activity in the brain, driven at least partly by reports that broadband activity can confound the estimation of narrowband, oscillatory activity (e.g., Cunningham et al., 2023; Donoghue et al., 2020a, 2022). In order to separate these components, however, researchers must make assumptions about how they combine to create the observed aggregate signal recorded from, for example, an EEG electrode or an MEG sensor.

Specifically, researchers must decide whether to model the spectrum as the *sum* or the *product* of broadband and oscillatory activity, with the additive approach assuming independence and the multiplicative approach assuming *inter*dependence between the two components. The difference between these two models has previously been addressed by us and others (e.g. Gyurkovics et al., 2021; Brake et al., 2024; Hu et al., 2024; Bloniasz & Stephen, 2026). However, with the expanding practice of employing separation approaches to estimate oscillatory power (or equivalently, amplitude) following the removal of broadband activity, the implications of different separation strategies warrant further discussion. Here, our goal is to provide a simple illustration of why it is necessary for researchers to make explicit their assumptions regarding additivity/multiplicativity of components, as these choices can have profound effects on the estimation of oscillatory activity and the resulting conclusions one might draw.

Broadband activity in neural data has a characteristic shape in the frequency domain: its power decreases as a function of frequency, following a power law of the form 1/*f*^x^, where *f* is frequency and *x* is the spectral exponent that defines how quickly power drops off as a function of frequency (e.g. Donoghue et al., 2020b; He et al., 2010; He, 2014, Miller et al., 2009). This 1/*f*-like shape reflects aperiodic neural activity, i.e., activity that does not repeat predictably over time (unlike oscillatory patterns). Traditionally, this aperiodic activity was assumed to be a stationary “background” that was typically ignored when analyzing oscillatory activity. However, recent observations challenge this view and clearly suggest that aperiodic activity changes in relation to cognitive processing (Cunningham et al., 2023; Gyurkovics et al., 2022; Kalamala et al., 2024; Lendner et al., 2023; Waschke et al., 2021), clinical status (Donoghue, 2025), and developmental stage and aging (Voytek et al., 2015; McSweeney et al., 2023; Turner et al., 2023; Clements et al., 2021; Stanyard et al., 2024; Warren et al., 2025). In research where oscillatory patterns, which manifest as narrowband peaks in the frequency domain, are the signals of interest, aperiodic activity can therefore represent a confounding variable that should be addressed prior to analysis.

Consequently, recent years have seen a proliferation of methods allowing for the separation of narrowband (potentially oscillatory) and broadband (non-oscillatory, aperiodic) signals (e.g. Brady & Bardouille, 2022; Clements et al., 2021; De Blasio & Barry, 2021; Demanuele et al., 2007; Donoghue et al., 2020a; Hu et al., 2024; Kałamała et al., 2026; Ouyang et al., 2020; Pascual-Marqui et al., 1988; Wen & Liu, 2016; Whitten et al., 2011; Wilson et al., 2022), with the *specparam* (spectral parameterization; formerly *fooof*) toolbox (Donoghue et al., 2020b), emerging as the most popular choice in the field. These methods, which seek to isolate oscillatory activity by removing the contribution of aperiodic activity, typically do not specify the assumed relationship between narrowband oscillatory and aperiodic activity (i.e., the assumption is often either implicit or left to the user). This is a problematic practice, as different assumptions necessitate different separation strategies, and as we demonstrate, these choices substantially impact measurement of the signals of interest.

Approaches to component separation rely on one of two fundamental models describing the relation between oscillatory and aperiodic activity (see Fig. 1). Multiplicative models assume that broadband and oscillatory activity multiply to create the observed spectrum. In these models, recovering the signal of interest (typically narrowband power assumed to reflect an oscillation) requires *dividing* the spectrum by an estimate of broadband activity. Additive models, on the other hand, posit that the components are independent and sum together linearly (e.g. Grandchamp & Delorme, 2011). To recover the former, an estimate of the latter needs to be *subtracted* from the spectrum. Which model is most appropriate to the underlying neurophysiology is not definitively known and plausibly situation dependent. However, available evidence that aperiodic and oscillatory activity are *not* correlated in the resting scalp EEG signal favors the additive model (Gyurkovics et al., 2022; Koenig & He, 2025) although evidence also exists that the multiplicative model might be more appropriate under certain circumstances (see e.g., Brake et al., 2024).

**Figure 1.**
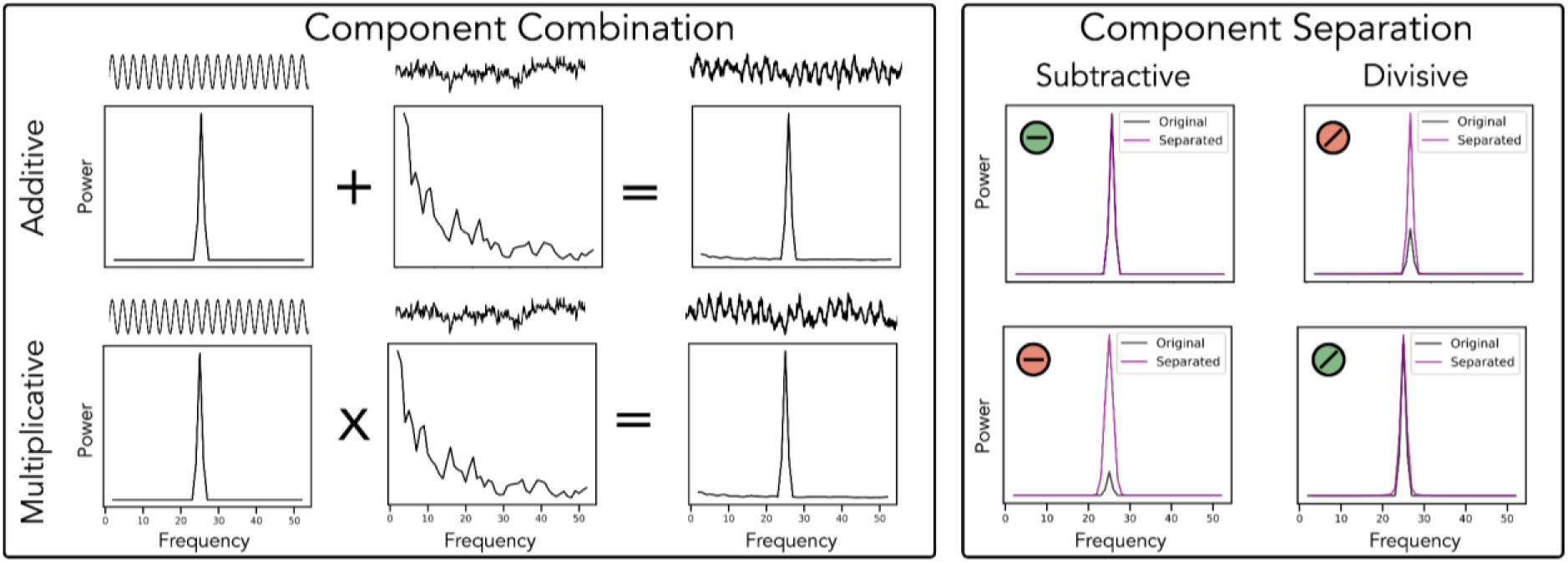
Schematic of component combination and separation. In the Component Combination panel, simulated time series (shown above their corresponding power spectra) are created for periodic (left) and aperiodic (middle) components, which are then combined together using either an additive (top row) or multiplicative (bottom row) approach to create combined time series (right). In the Component Separation panel, the combined power spectra are then analysed to isolate peaks using either subtractive (linear-subtraction) or divisive (linear division or equivalently log-subtraction) removal of the aperiodic component. In doing so, applying the peak separation approach that corresponds to the signal creation (indicated by green shading of ‘-’ or ‘/’ symbols, reflecting subtractive and divisive approaches, respectively) successfully isolates the initial components, whereby the recovered spectral peak (purple) matches the original component (black). Applying a separation discordant with how the signals were combined (indicated by red shading) leads to mis-estimations of the input components.

Given that existing signal decomposition techniques vary regarding which of the two models is assumed, often without making this assumption explicit, researchers may find themselves in the situation of intending to apply an additive model of neural signals, but inadvertently performing multiplicative computations, or vice versa. This issue (a mismatch between assumed additivity/multiplicativity of the source as compared to the separation approach) is likely exacerbated by the fact that neural power spectra are often represented in log-space (i.e., power or amplitude values are log transformed), as subtraction in log-space is equivalent to division in linear space (i.e., log_10_(a) - log_10_(b) = log_10_(a/b)). The net result can be an inadvertent misalignment between applied measurement and intended interpretation of the oscillatory signal.

In this paper, we argue that when spectral separation is performed, it is vital for researchers to explicitly justify which of these assumptions (additive/multiplicative) is endorsed and select the appropriate separation strategy, because a mismatch between assumption and strategy can lead to a serious distortion of the to-be-recovered signal (see Fig. 1). In a similar vein, we recently showed that when a divisive baseline normalization strategy was used to emphasize post-stimulus activity in time-frequency analyses, completely illusory post-stimulus dynamics could emerge (Gyurkovics et al., 2021). This finding then informed consensus guidelines on the reporting of time-frequency domain analyses of neural time series (Keil et al., 2022).

Because accurate signal separation is impossible without first specifying assumptions regarding signal combination, the considerations we discuss herein are relevant for all toolboxes and algorithms that aim to separate broadband and narrowband signals. In the present paper, the analyses employ the *specparam* (formerly *fooof*) package (Donoghue et al., 2020b), in which the power spectrum is parameterized as the combination of an aperiodic component and a set of overlying peaks reflecting frequency-specific (putatively oscillatory) activity. By default, however, aperiodic-adjusted peak power estimates from *specparam* have (until recently) been derived from subtraction of log-transformed power spectra, which assumes multiplicative signal generation. We illustrate in simulation that if the data is additively combined, divisive separation can distort narrowband power estimates, leading to over- or under-estimation. We also show in real empirical datasets, including resting-state and task-related data, that the choice of divisive vs. subtractive signal separation alters interpretations, further emphasising that researchers should consider and explicitly report these analytic choices. To support this, the *specparam* method (as of version 2.0; https://specparam-tools.github.io/) has been updated to support explicit selection of whether to model and report peak power estimates as additively or multiplicatively extracted from the aperiodic component, allowing users to choose, compare, and report these choices.

## 2. Results

### 2.1. Simulations

#### 2.1.1. Separation strategy impacts aperiodic-adjusted power estimates

To first demonstrate the effects of the separation approach, we created several simple simulations using either additive or multiplicative combination of periodic and aperiodic components, followed by subtractive or divisive peak separation. The associated figures (Figs. 1-2) show the impact of applying peak separation approaches consistent or inconsistent with how the components were combined. While correct specification accurately reconstructs simulated peak heights, incorrect specification does not (Fig. 1). Notably, these misestimates due to model misspecification can lead to a variety of reported findings (Fig. 2, rightmost column). Driven by differences in exponent and/or peak height values and depending on the ‘rotation point’ frequency (the frequency at which spectra cross-over for a given exponent difference), model misspecification can lead to both false positive and false negative results, with either increases or decreases in peak amplitude.

**Figure 2.**
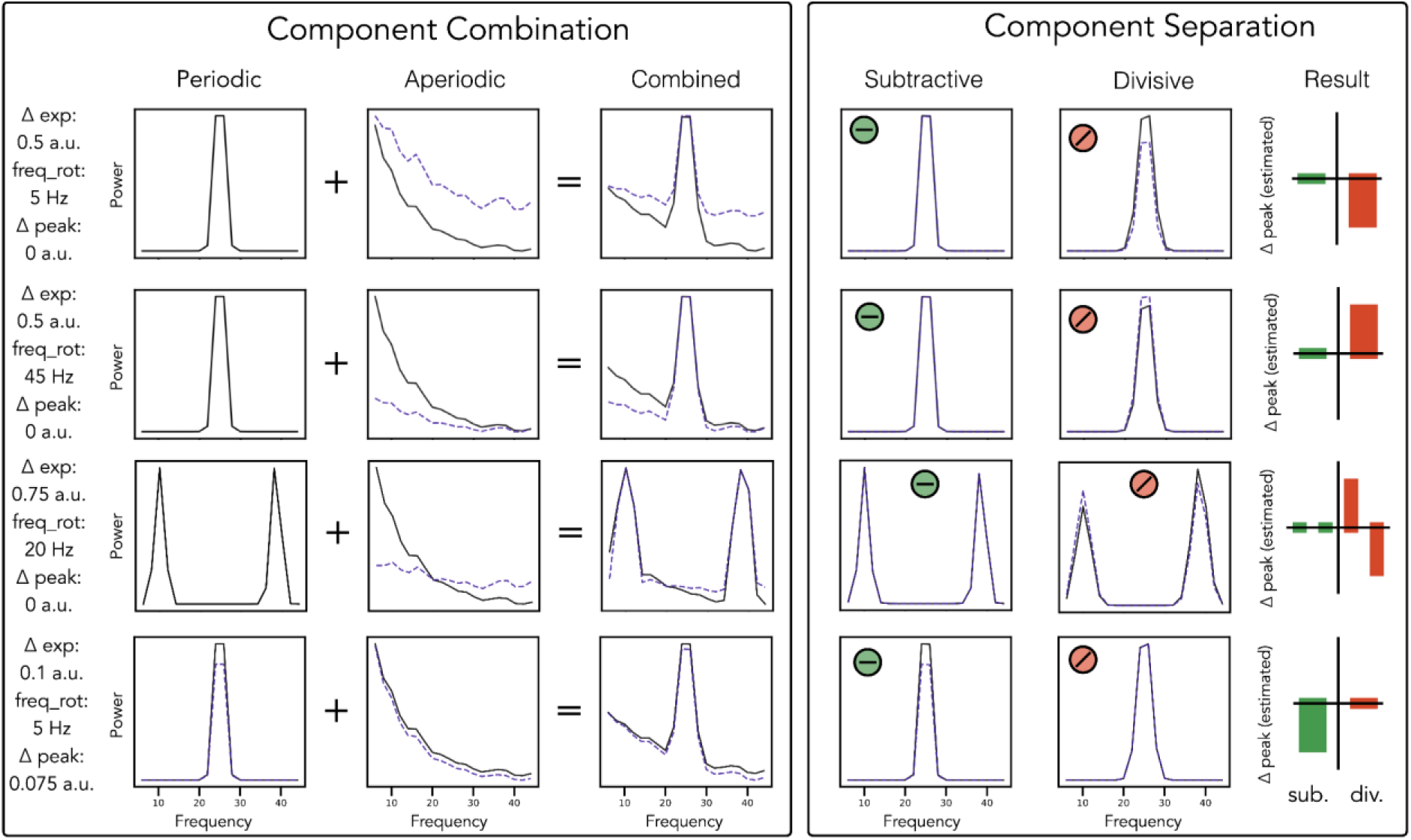
Examples of correct and incorrect estimations based on alignment of combination and separation approach. In the Component Combination panel, periodic and aperiodic components are created separately and combined additively to create two different conditions. In each row, the parameter description (left) describes the difference in the 2nd condition (dashed purple) as compared to the 1st (black). In the Component Separation panel, subtractive or divisive peak extraction techniques are applied to estimate the difference in peak power between conditions. Across these examples, with are combined additively, a subtractive separation (correct specification, indicated by shaded green minus symbol) correctly estimates the simulated changes, whereas various kinds of mis-estimations can arise if using a divisive separation (incorrect specification, indicated by shaded red division symbol). This includes false positive peak decreases (row 1) or peak increases (row 2) where the correct estimation is no difference in peak height. In situations with multiple peaks, false positive and negative peak deviations can co-occur, with measured differences depending on the relationship between peak frequency and exponent rotation frequency (row 3). Finally, true peak differences can be mis-estimated, for example a false negative finding of no difference when the correct estimation is a peak decrease, which is masked by the aperiodic change due to the mis-specification (row 4). Note that these simulations are curated to show specific example cases, using specified parameters for exponent differences, exponent rotation points, and peak height differences.

Following these initial simulations, we next sought to systematically evaluate the impact of divisive vs. subtractive separation on estimates of narrowband power derived from simulated spectra in which aperiodic components with varying offset and exponent values were additively combined with narrowband oscillatory activity. Simulated spectra were generated by adding a burst of 12 Hz activity (periodic ‘alpha’ signal) to aperiodic broadband activity. The parameters of the simulated aperiodic activity were modulated in two runs. In the first run, the offset value was kept constant while the exponent value increased across 20 levels; in the second run, the exponent was varied and the offset held constant. These two sets of simulations were conducted 3 times, increasing the amplitude of the alpha signal in each instance (modelling an increase in signal-to-noise ratio). Importantly, alpha amplitude was fixed across the 20 levels *within* each run. For each spectrum, we quantified the magnitude of the alpha signal in two ways: after divisive separation of aperiodic and periodic activity (i.e., by using the aperiodic parameters output by *specparam* to recreate the broadband part of the spectrum, and then subtracting this from the full spectrum in log space which closely approximates the periodic parameters output by *specparam)*, and after subtractive separation (i.e., by using the aperiodic parameters output by *specparam* to recreate the broadband part of the spectrum, and then subtracting this from the full spectrum in linear space).

Figure 3 shows aperiodic-adjusted periodic power estimates in the alpha range as a function of a range of exponent (Fig. 3A) or offset values (Fig. 3D) in simulated spectra. Regardless of which aperiodic parameter was varied, the results suggest that as the proportion of power attributable to broadband background activity increased in the alpha range (i.e., as the offset increased or the spectrum rotated around a fulcrum outside the alpha range), the aperiodic-corrected alpha power values extracted using divisive correction (i.e., assuming an–*incorrect*–multiplicative model) decreased, at all investigated signal-to-noise ratios (Fig. 3B, E). This pattern emerged despite the fact that the actual oscillatory burst added to the background noise was identical in *all* conditions. As expected, when the correction for aperiodic activity was done by subtracting non-log-transformed aperiodic values from non-log-transformed full spectra, alpha power values did not change as a function of either exponent (Fig. 3C) or offset (Fig. 3F).

**Figure 3.**
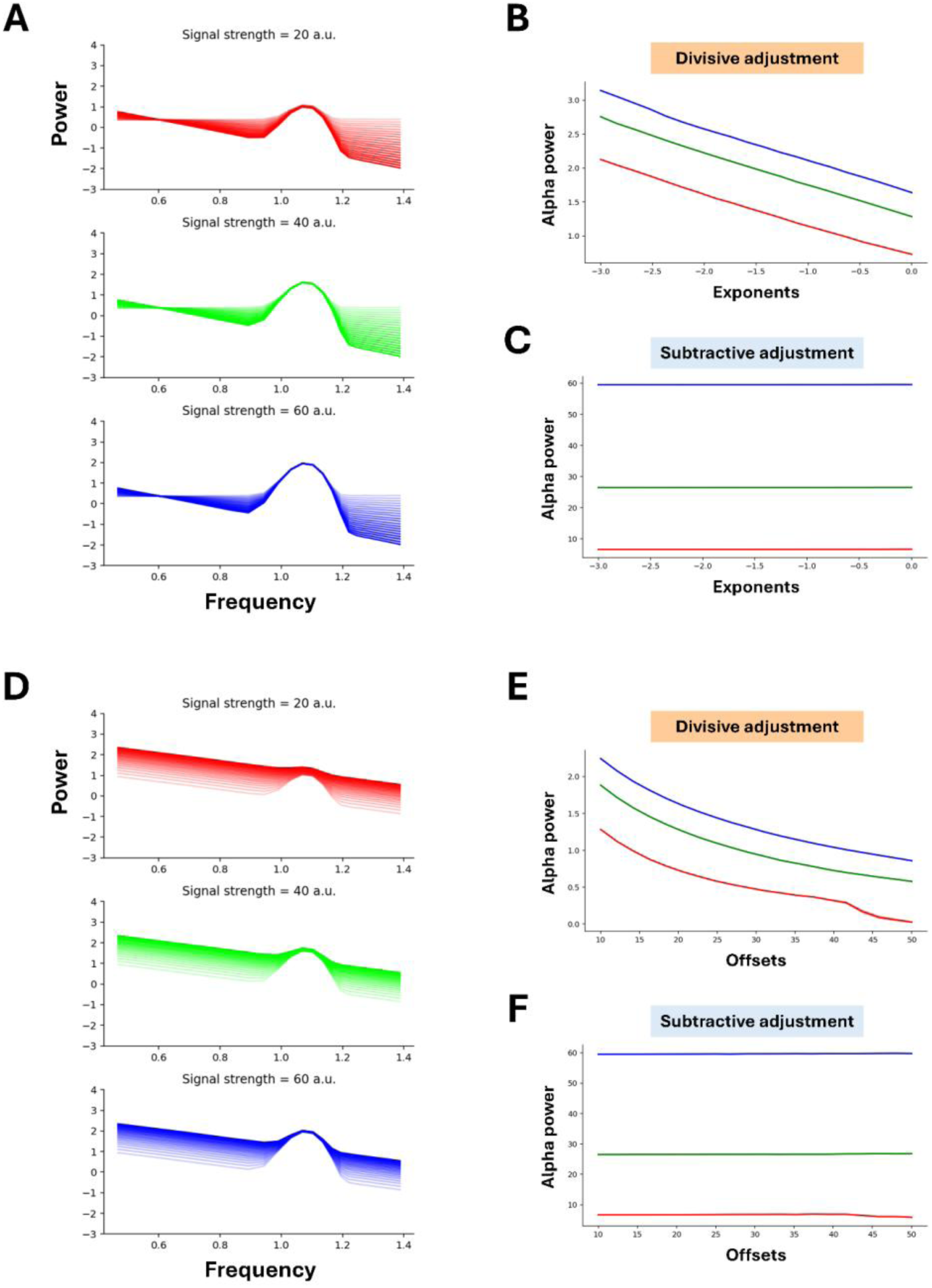
Aperiodic and aperiodic-adjusted periodic parameters as a function of the simulated aperiodic parameter. **A**) Average simulated (log-transformed) spectra as a function of signal-to-noise ratio (subpanels) and the change in the exponent (color transparency). The more opaque the color, the larger the absolute value of the simulated parameter. The magnitude of the alpha signal was fixed across all conditions. **B**) Periodic power in the alpha after aperiodic correction in log space, as a function of exponent values. This is an example of divisive aperiodic correction, and mimics the steps specparam follows for aperiodic adjustment. **C**) Periodic power in the alpha range after aperiodic correction in the linear space. This constitutes subtractive correction. **D**) Same as A, but with offsets varied. **E**) Same as B, but as a function of offsets. **F**) Same as C, but as a function of offsets.

#### 2.1.2. Separation strategy can lead to the misestimation of oscillatory power differences

Next, we investigated the impact of separation strategy on the estimation of the effect size of an oscillatory difference between conditions (using additive signal combination as ground truth). Spectra were simulated in two conditions within two runs. In the first run, offset was held constant across conditions, but exponent was allowed to differ between conditions, with the difference increasing across 21 levels. In the second run, the exponent was held constant across conditions, and offset changed (for 21 levels of difference in offset between conditions 1 and 2). The two conditions also differed in the amplitude of the alpha signal added to the broadband component. The alpha amplitude in each condition—and therefore the amplitude difference between conditions—was held constant across all runs and iterations. We then quantified the difference in estimated alpha power between conditions for each spectrum, once after divisive separation, and once after subtractive separation, as above.

Fig. 4 presents the results of the simulation runs exploring a range of conditional difference values for the exponent (Fig. 4A) and the offset (4B), while keeping the oscillatory alpha difference constant, and thus provides a systematic investigation of the phenomenon shown in Fig. 1. Notably, when the separation approach matched the known generative model (additive), the estimated oscillatory effect was relatively constant regardless of aperiodic variation (blue lines in Fig 4). When the correction approach did not match the generative model (i.e., divisive correction was used), the estimated oscillatory effect systematically differed with the size of the aperiodic difference between conditions.

**Figure 4.**
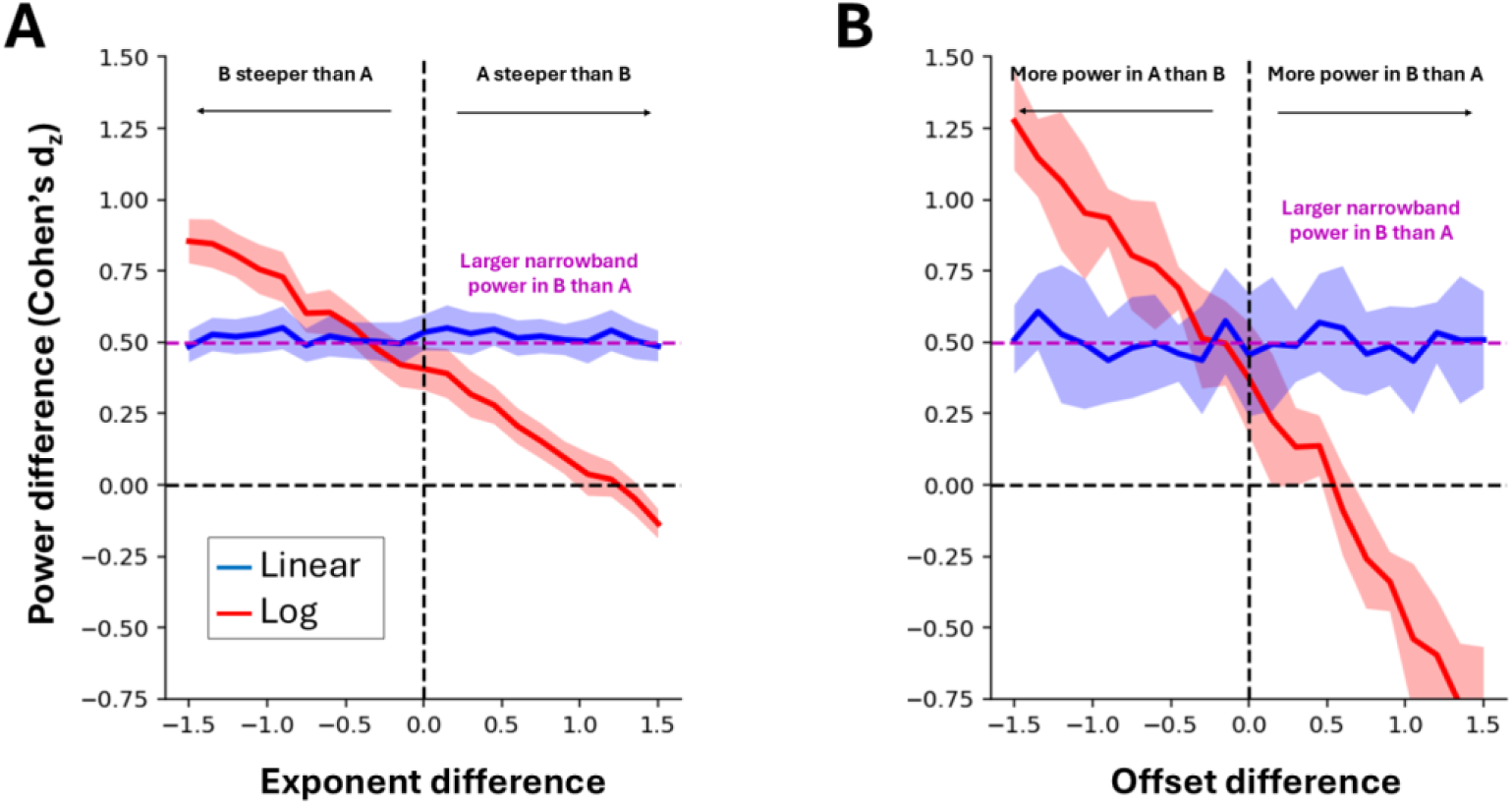
The effect of divisive and subtractive separation strategies on estimation of oscillatory effects when aperiodic effects are also present and the ground truth is additive. **A**) Simulation of effects of divisive vs. additive separation on estimation of differences in oscillatory power between two conditions with different aperiodic exponents and different alpha peaks (over a range of exponent difference values). The effect size of the power difference in alpha was measured after flattening the mean spectra in linear space, and in log space. The ground truth alpha difference between conditions was fixed (B > A, see text in magenta) across exponent difference levels, and is indicated by the dashed magenta line. The Y axis shows the effect size (Cohen’s d_z_) for the exponent difference. Shading around curves indicates 95% confidence interval. **B**) Same as B, but for offset differences.

#### 2.1.3. Separation strategy affects conclusions in real data

We next sought to examine the impact of separation strategy in analyses of real data. We first examined age-related changes in alpha power in a resting state data set (Langer et al., 2017), and show that results change as a function of peak separation approach. In an empirical dataset of eyes-closed resting EEG data (n=111), we observed an expected posterior distribution of peak alpha power (Figure 5A). Focusing on posterior electrode E72 (approximately Pz), we recomputed peak power using both subtractive and divisive aperiodic separation (Fig. 5B). We then examined the relationship between isolated peak alpha power (after linear or log subtraction) and age (Fig. 5C). We found that linearly-subtracted alpha power was significantly correlated with age (*r*=-0.32, CIs: [−0.48- −0.15], *p*<0.01), whereas log-subtracted alpha power was not (*r*=0.14, CIs: [−0.06 – 0.32], *p*=0.15). A bootstrapped test for a difference between these correlations was significant (Δ*r*=-0.46, CIs: [−0.58 - −0.34], *p*<0.01). This demonstrates that the choice of correction strategy can significantly impact the apparent relation between estimated spectral features and demographic features such as age. Notably, the estimated difference between measured correlations is large, showing the different approaches can lead to quite different conclusions.

**Figure 5.**
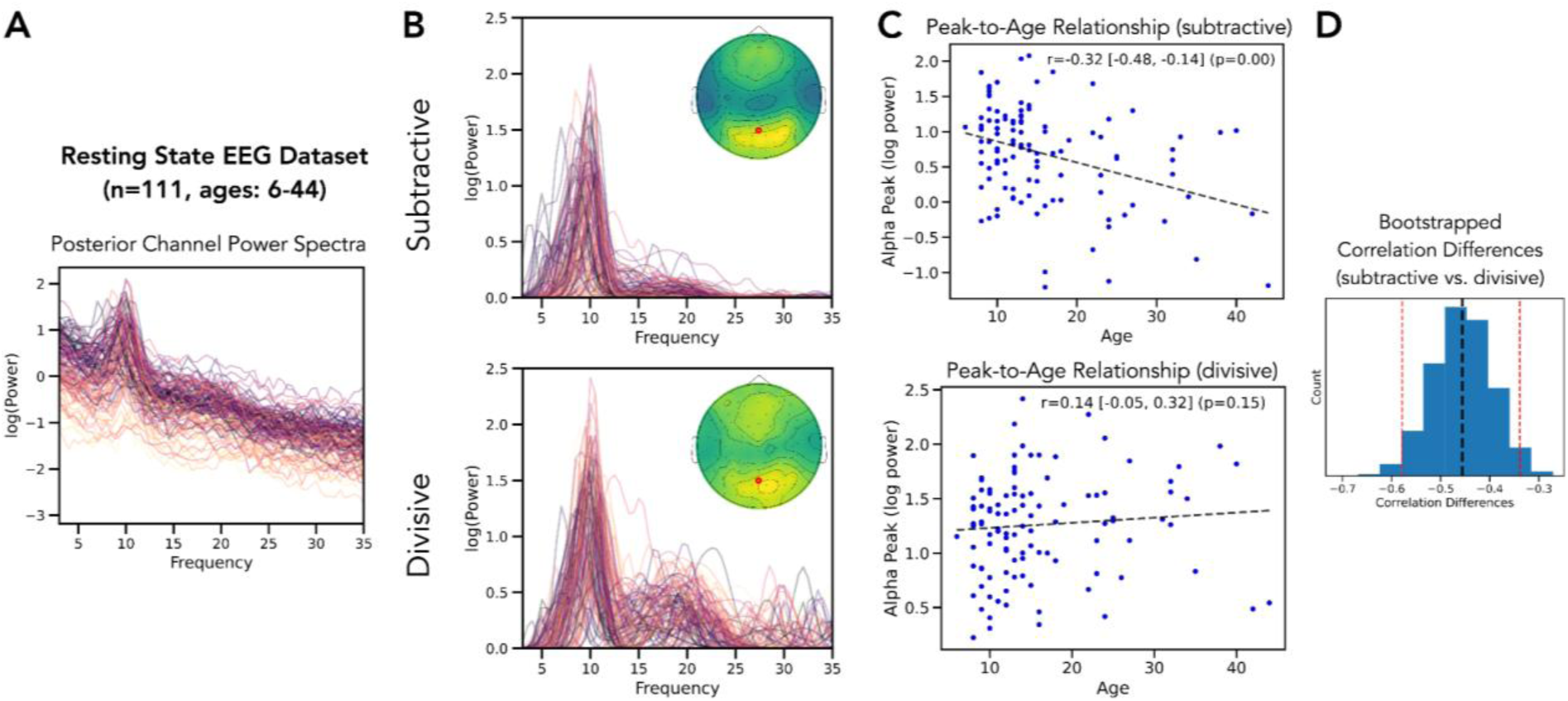
Alpha-power/age relationship as a function of periodic-aperiodic separation approach in empirical resting state EEG data. A) Example power spectra (channel E72, approximately Pz) of resting-state EEG data (n=111; MIPDB dataset from the ChildMind Institute). Power spectra are colored by age (age range=6-44; younger lighter). B) Analysis of periodic activity using a subtractive approach (linear-subtraction; top) or divisive approach (linear division or log-subtraction; bottom) showing the isolated periodic components for an example channel. Insets show the topography of alpha power with the selected electrode indicated. C) Analysis of the relation between peak power in alpha range (6-14 Hz) and age, separately for peak power computed with the subtractive (top) or divisive (bottom) approach. Inset text reports the Spearman correlation rho-value, associated p-value and bootstrapped confidence intervals (see Methods). D) Bootstrapped correlation difference, in which resampling is used to estimate the difference between the correlation magnitudes using each approach, showing a significant difference between correlations computed with subtractive vs. divisive peak estimations (delta r=-0.46, CI: [−0.58--0.34], p<0.01). The black dashed line indicates the mean correlation difference value of the distribution of bootstrapped estimates, and the red dashed lines indicate the 95% confidence interval.

Finally, we examined the impact of separation strategy in real data in an analysis of task-related activity, using data from Cunningham et al. (2023). In this study, among other questions, the authors examined the effects of correcting for aperiodic activity on the relation between pre-stimulus alpha-band (8-12 Hz) power and visual awareness. In the literature, it has been extensively reported that increases in ongoing posterior alpha-band power tend to precede failures of subjective visual awareness (for a review and synthesis, see Samaha et al., 2020). Cunningham et al. (2023) observed that (a) systematic shifts in the aperiodic component of the spectrum also predicted visual awareness, and (b) after separation of aperiodic and oscillatory components of the spectrum through subtraction (assuming additive generation), the relation between alpha power and reports of visual awareness was still present, though reduced in magnitude (reproduced in Fig. 6B; compare with Cunningham et al., 2023, Figures 6F-G). This led to the conclusion that both features of the signal together produce the nominally alpha-band effects often reported in the literature. Here, we illustrate how the conclusions would differ had Cunningham et al. (2023) instead assumed a multiplicative model (subtracting the aperiodic component from the log-transformed power spectrum). We refer to alpha power before any correction for aperiodic activity as *unadjusted* alpha power and after any correction for aperiodic activity as *adjusted* alpha power.

**Figure 6.**
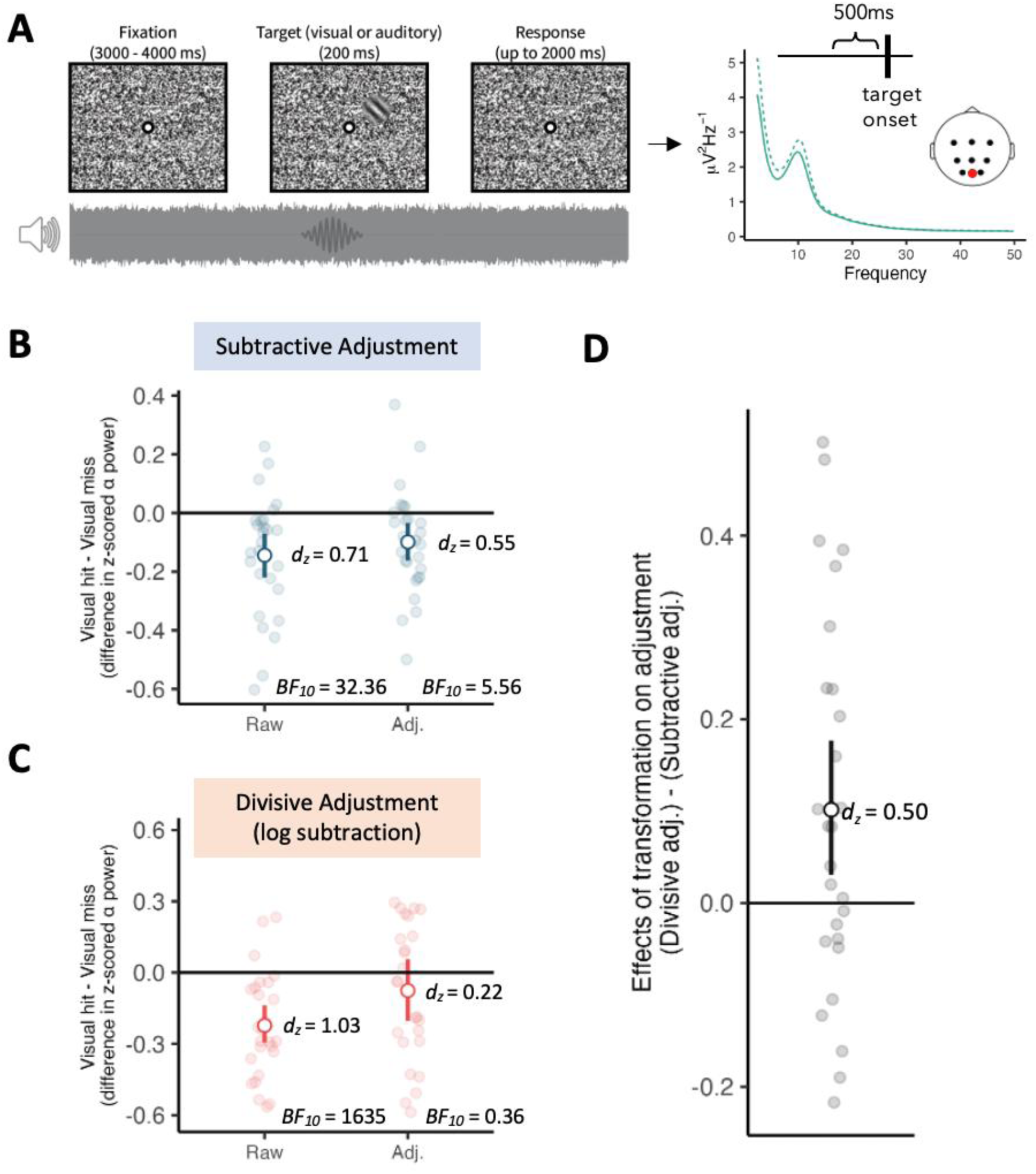
Reanalysis of data from Cunningham et al. (2023). **A)** (Left) Task schematic. Participants had to detect near-threshold auditory or visual targets embedded in auditory or visual noise. (Center) Time window of analysis and the electrode of interest. (Right) Average pre-stimulus spectra for trials where participants detected the target (hits) and trials where they did not (misses). **B)** Results as in Cunningham et al. (2023): The difference in alpha power between hits and misses, before (“Raw,” for unadjusted) and after (“Adj.”, for adjusted) the removal of aperiodic “background” activity through linear subtraction. d_z_ = Cohen’s d_z_ for one-sample or paired-samples comparisons; BF_10_ = Bayes Factors, where larger values indicate more support for the alternative hypothesis over the null. Points and interval estimates are means and 95% bootstrapped CIs. Dots represent individual data. **C)** Same as B but with adjustment in log space (i.e., divisive separation). **D)** The difference between the effect after divisive adjustment (“Adj.” in panel C) and the effect after subtractive adjustment (“Adj.” in panel B).

In Fig. 6, the difference values for the contrast of ‘visual hit - visual miss’ are plotted for both adjusted and unadjusted alpha power assuming an additive model (Fig. 6B) and a multiplicative model (i.e., subtraction after log-transformation; Fig. 6C) for representative electrode Oz. Across individuals, a divisive approach (subtractive correction in log-space) exaggerated the consequences of signal adjustment compared to a subtractive approach, *t*(26) = 2.58, *p* = .016, *d_z_* = 0.50 (Fig. 3D). As in the original paper, the effect of aperiodic correction in linear space (additive model) was to reduce, but not eliminate, the narrowband oscillatory effect (reducing the associated Bayes Factor from *BF_10_* = 32.36 in the uncorrected data to *BF_10_* = 5.56 in the aperiodic-adjusted data). However, the consequences of a divisive approach were more dramatic, effectively eliminating the narrowband effect: it reduced the associated Bayes Factor from *BF_10_* = 1645 in the uncorrected (log-transformed) data to *BF_10_* = 0.36 in the aperiodic-adjusted data, a drop from substantial evidence favoring a difference between conditions to weak evidence favoring no effect.

## 3. Discussion

Previous studies have demonstrated that broadband, aperiodic activity can confound estimates of narrowband oscillatory activity in brain signals (e.g., Donoghue et al., 2020; Voytek & Knight, 2015), leading to the argument that it is critical to separate these components for accurate measurement. To do this, researchers must have a clear model of the relation between the two components because different models necessitate different component separation strategies. Here, we demonstrate that a mismatch between model and measurement strategy can have serious consequences for the results of oscillatory analyses and, in turn, the interpretation of findings.

We considered two models of signal combination: an additive model where oscillatory and aperiodic activity are independent, and a multiplicative model where oscillatory and aperiodic activity scale together (and are therefore dependent). The former model requires subtraction to separate the components, and the latter requires division (or subtraction in log space, which is an equivalent approach). Our results show that if one endorses an additive model, techniques that implicitly or explicitly involve division (as has been the default in the *specparam* toolbox; Donoghue et al. 2020) will lead to distorted oscillatory estimates. To address this concern, which model to use can now be explicitly specified in *specparam*.

Misspecification can lead to both under- and over-estimations of oscillatory changes. In simulations which systematically increased the contribution of aperiodic power to the narrowband frequency range of interest, divisive separation strategies led to increasing underestimation of narrowband power. This pattern emerged regardless of whether the aperiodic power increase was due to a rotational shift in the spectrum (a change in the exponent) or to a broadband shift (a change in the offset). As expected, when the generative model of the data and the separation strategy matched (i.e., subtractive correction was used for additively generated data), power estimates for oscillatory activity remained constant around the ground truth value regardless of aperiodic changes. These findings are also in line with our previous simulations in Cunningham et al. (2023) and Gyurkovics et al. (2021), as well as with the findings of Brake et al. (2024, see e.g., Fig. 5).

Importantly, inappropriate divisive separation of additively generated signals can also obscure or even misrepresent narrowband differences between conditions. We simulated a scenario where conditional differences in aperiodic and oscillatory activity co-occurred but were independent and found that subtractive separation (i.e., the correction that matched the generative process in the simulation) was largely insensitive to the size of the aperiodic difference between conditions, yielding accurate recovery of oscillatory effects. Divisive correction, on the other hand, led to an *overestimation* of the oscillatory effect when the condition with higher narrowband power was also the condition with lower broadband (aperiodic) power; but an *underestimation*, or even a complete misestimation of the sign of the effect when that condition was associated with higher aperiodic power, and was thus subject to the same shrinking effect described above. Practically, this means that researchers risk concluding that one condition has higher narrowband power in a given frequency range compared to another after correcting for aperiodic activity, even if in reality the opposite is true, depending on the separation choices employed, and the effect sizes of the oscillatory and/or broadband changes in the data.

Through our reanalyses of real datasets, we also provide real-world examples of how divisive and subtractive correction approaches can yield different conclusions. In a developmental dataset (Langer et al, 2017), we find a difference in whether an age-related change in alpha power is detected depending on which separation approach is taken. Similarly, in the re-analysis of the detection task of Cunningham et al. (2023), the well-established pre-stimulus alpha power difference between hits and misses (see e.g., Samaha et al., 2020) disappeared after correcting for aperiodic activity using a divisive approach. Faced with these results, one might conclude that the original ‘alpha-band’ effect was entirely or nearly entirely attributable to the broadband component of the signal. Subtractive correction, however, suggested two independent broadband/narrowband pre-stimulus effects. Relative to our simulations, these results would fit on the right-hand side of Fig. 4A-B (and the fourth row of Fig. 2), where the red line (for divisive separation) crosses 0 on the x-axis (corresponding to no oscillatory effect after divisive correction), and the blue line (for separation in linear space) is close to the ground truth effect, indicating the co-occurrence of narrowband and broadband effects. Naturally, we do not have access to ground truth in these examples and thus cannot adjudicate between separation strategies, but Cunningham et al. (2023) do note that after subtractive separation, the correlation between aperiodic and periodic estimates was limited, consistent with an additive model.

These data also highlight that the scenario we simulate—in which condition differences are present along both aperiodic and narrowband dimensions—is not unrealistic (for further evidence, see e.g., Gyurkovics et al., 2022; Kałamała et al., 2022; Waschke et al., 2021). Even more importantly, the two conditions simulated here could also correspond to two populations that differ in aperiodic signal structure, such as younger vs older adults (e.g., Clements et al., 2021; Dave et al., 2018; Turner et al., 2023; Voytek et al., 2015), or individuals with or without certain clinical disorders (e.g., Dakwar-Kawar et al., 2024; Robertson et al., 2019, see Donoghue, 2025 for a review). In these situations, misestimation of narrowband differences between groups is a realistic concern if divisive correction is performed but an additive generative model is assumed, or vice versa.

The present paper demonstrates that choosing an additive or multiplicative separation strategy, even implicitly, can substantially affect subsequent analyses and conclusions. Thus, the work presented here also highlights the need for new research to understand when and under what circumstances the underlying signal generation and source combination is multiplicative or additive. Thus far, existing evidence seems to favor the additive model in most circumstances (e.g., Brake et al., 2024; Gyurkovics et al., 2021; Koenig & He, 2025, Bloniasz & Stephen, 2026), though there has been some discussion of alpha dynamics reflecting filtered noise (Gebber et al, 1999) or a damped oscillator (Evertz et al, 2022), which would suggest different kinds of modeling strategies that could include multiplicative combinations.

In conclusion, through simulations and real data, we highlight a critical, though largely overlooked issue in the literature on narrowband and broadband activity. Namely, that researchers often do not specify what relationship they assume to exist between these two types of activity. This assumption has fundamental implications for how broadband, aperiodic activity should be removed from the power spectrum to yield isolated estimates of narrowband oscillatory power. We illustrate the importance of considering this assumption through the use of *specparam*, an influential toolbox in the field, but it is critical to note the implications of our findings are not unique to this toolbox and apply to any correction of frequency-specific activity relative to a broadband component. Therefore, this assumption must be considered and made explicit regardless of the algorithm used for signal separation and parameterization. We recommend that authors clearly state what method they used to obtain aperiodic-corrected narrowband estimates of power from the raw power spectrum: subtraction or division (alternatively, subtraction in log space), and why they endorse the corresponding assumption regarding signal generation: additive or multiplicative generation, respectively. This is in line with recent recommendations for baseline normalisation in time-frequency analysis (Keil et al., 2022). To support these additional steps, a choice of separation can now be explicitly specified in the *specparam* toolbox (version >= 2.0). Future work should further evaluate these choices, including in connection to modeling of signal sources, and could also consider more complex forms of signal separation with novel methods and models. Regardless of the specific aims of any given study, we advocate for explicit reporting of both the method of separation and the underlying rationale.

## 4. Methods

### 4.1. Procedure

#### 4.1.1. Simulations

Simulations were done using Python (v3.13), including the *neurodsp* Python toolbox for time series simulations and signal processing (Cole et al., 2019). Time series were simulated as separate periodic and aperiodic components, which could be combined following additive or multiplicative approaches. In the time domain, the additive approach is the sum of the two components. For the multiplicative model, the combination is the convolution between the two components, since multiplication in the frequency domain is equivalent to convolution in the time domain. Periodic components were simulated as continuous power, ongoing oscillations (unless otherwise noted) which were scaled to different peak-to-peak heights to manipulate oscillatory power, and aperiodic components were simulated as colored noise signals to specified 1/*f* exponent values by spectrally rotating white noise (Timmer & Konig, 1995). Simulations were created at a default sampling rate of 500 Hz.

For the initial simulations shown in Figures 1 & 2, segments of length 30s were generated, modelling a resting state recording. Default parameters used a peak center frequency of 25 Hz for single peaks (10 and 40 for two peaks) and an exponent value of −1. For comparisons between spectra, secondary spectra were created using specified differences of height and exponent, with a given exponent rotation point (i.e., the frequency where two spectra of different exponents cross over), as specified for each individual simulation. These paired spectra were then analyzed by removing the simulated aperiodic component using a subtractive or divisive approach, with the isolated periodic components then compared to the originally simulated periodic components and examined for accuracy of the reconstruction. Note that these simulated differences are curated example parameters chosen to highlight potential differences, serving as exemplary ‘point estimates’ within the continuous variation in outputs that is observed when continuously varying input parameters.

Next, we further investigated the effect of aperiodic parameter changes on narrowband oscillatory power estimates separately for subtractive separation (subtraction in linear space) and divisive separation of components (subtraction in log-space). Note that we use “narrowband”, “oscillatory”, and “periodic” interchangeably in this report because, in our simulated data, narrowband activity was always oscillatory/periodic in nature. For each of 20 aperiodic parameter values, one hundred epochs (1024 ms each) were generated (exponents were varied in one run, offsets in another). These shorter epochs model a task-based scenario where more transient, stimulus-related changes are of interest to the researchers. In the exponent run, exponent values consisted of 20 values ranging from −3 to 0, with the offset fixed at 10 (arbitrary units). The rotational point of the spectra was at 4 Hz. In the offset run, offsets consisted of 20 values ranging from 10 to 50, with the exponent fixed at −2.

For all epochs, the aperiodic component was simulated using *neurodsp* (as above). An 8-cycle-long burst of ∼12 Hz activity (created by multiplying a ∼12 Hz sine wave and a Gaussian taper; approximating alpha-rhythmic activity) was then generated in the time domain and converted to the frequency domain using the Fourier transform as implemented in the fft.fft function in Python 3.13. This component was then added to the spectrum of the aperiodic component, consistent with additive signal generation. We use the additive model as an example because, given the complications of log-transformation, it is easier to inadvertently perform divisive correction when one’s assumptions are additive than vice versa. For each of the 20 aperiodic parameter levels, three conditions were simulated, corresponding to three different signal-to-noise ratios, by multiplying the ‘alpha’ signal with weights of 20, 40, and 60 (arbitrary units). Epochs were subsequently averaged in the frequency domain. Then, the 2-25 Hz frequency range in each resultant spectrum was decomposed using *specparam* with the following parameter settings: *peak_width_limits*=[2,8], *max_n_peaks*=1, *min_peak_height*=0.3, *peak_threshold*=2.0, *aperiodic_mode*=’fixed’.

To illustrate the effects of divisive separation, the aperiodic parameters (offset and exponent) produced by *specparam* were used to generate the aperiodic model component which was then subtracted from the full spectrum in log space. Power values were then averaged between 7-13 Hz in the subtracted spectrum. This resulted in power estimates largely identical to using the peak height metric output directly by *specparam* (average correlations between the two metrics ranged between 0.97-0.99 across SNR levels). To illustrate the effects of subtractive separation, the same steps were followed but separation was done without log transformation, i.e., in linear space. For both the offset and the exponent runs, all 20 × 3 simulations were run 250 times, and results were averaged across iterations.

Next, we examined the impact of inappropriate spectral parameterization on the measurement of true oscillatory group differences when aperiodic differences were also present between conditions (assuming an additive model of signal generation). To create aperiodic condition differences of varying magnitude, 21 parameter difference levels were simulated between two conditions, once for exponents, and once for offsets. In the exponent run, the exponent was fixed at −1 in one condition and ranged from −1± 0.15 in the other. The offset was fixed at 5 for both conditions. In the offset run, the offset was fixed at 5 for one condition and ranged from 5 ± 1.5 in the other. The exponent was fixed at −1 for both conditions. For each conditional difference level, 100 spectra were generated for each condition for 40 simulated subjects, and the power difference in the alpha range was measured after flattening the mean spectra in linear or log space. In both cases, the aperiodic component of the spectrum was estimated based on the values output by *specparam*, but it was subtracted from the full spectrum without log-transformation in the additive case, and after log-transformation in the multiplicative case. The ground truth ‘alpha’ difference between conditions was fixed for all parameter difference levels at Cohen’s *d*_z_ ∼ 0.5. 10 iterations of the simulation were run at each difference level.

#### 4.1.2 Empirical data

##### MIPDB dataset

To examine the impact of additive vs. multiplicative assumptions in empirical data analysis, we first used the openly available Multimodal Resource for Studying Information Processing in the Developing Brain (MIPDB) dataset of human EEG data from the ChildMind Institute (Langer et al., 2017). EEG analyses were done using Python (v3.13), including using the MNE toolbox (Gramfort et al., 2014). The dataset was accessed and processed as described in a previous investigation (Donoghue, Dominguez, et al., 2020). The original sample included 126 participants (aged 6–44 years; mean ± SD: 15.79 ± 8.03 years; 69 males), whose EEG was recorded using a 128-channel Geodesic HydroCel system. Of these, 111 had a complete 30-second segment from the first eyes-closed resting-state block and were included in the present analysis. Power spectra were calculated for each channel using Welch’s method, with 2-second windows and 25% overlap (Welch, 1967), giving 0.5 Hz frequency resolution. Power spectra were parameterized with *specparam* (formerly *fooof*; version 2.0.0rc7; Donoghue, Haller, et al., 2020), with the default fitting algorithm over the frequency range 3-35 Hz using a 1/f (‘fixed’) aperiodic fit function and a Gaussian peak function, with additional fitting parameters set as: peak_width_limits: (1-6 Hz), max_n_peaks: 6, min_peak_height: 0.1, peak_threshold: 2.0. Spectral models had an overall model r-squared of 0.97 across subjects and channels (as examined in the topographies), indicating good fits overall. For the main analysis, which examined channel E72 in the Hydrocel-129 montage (approximately Pz position), the average R-squared was 0.98, indicating good model fits.

Peak alpha power was extracted as the highest power peak between the range of 6-14 Hz, using a broad range due to potential variation in peak frequencies across the varied age-range of the dataset. This was done both with linear power subtraction (subtractive) and log power subtraction (divisive) to estimate both measures of alpha power. To examine the relationship between age and spectral measures, Spearman correlations were computed. Bootstrapping procedures were applied to compute confidence intervals for each reported measure and to test for differences between correlation magnitudes by testing whether resampled correlation differences are significantly different from zero (Wilcox, 2016), each using 5000 resamples with replacement.

##### Cunningham et al. dataset

We also used empirical data from Cunningham et al. (2023) to provide a practical illustration of the impact of assuming an additive vs. multiplicative model of signal generation in a task context (associated data available at osf.io/2u4g9). This study examined the impact of aperiodic signal separation on measurement of pre-target alpha-band power during visual detection (building on observations that increased posterior alpha-band power tends to precede failures of subjective visual awareness). For the purposes of the current paper, we focus on the contrast of within-individual differences in alpha power prior to visual hits relative to misses, using this contrast to illustrate how conclusions would change assuming an additive vs. multiplicative model.

The data are derived from those used to produce Figures 2F-G (and 4B/4D) of Cunningham et al. (2023). Interested readers are referred to the original paper for full details of preprocessing, fitting, and decomposition. Power in the alpha band was computed as the area under the curve in the range ± 2 Hz surrounding peak alpha frequency (for comparison with the original report). This metric was extracted both pre- and post-subtraction of the aperiodic portion of the spectrum (described as ‘unadjusted’ and ‘adjusted’, respectively), for both log-transformed and untransformed spectra. To facilitate comparison of log-transformed and untransformed outcomes, power values were standardized (*z*-scored) across individuals prior to computing contrasts. In addition to computing the effect sizes associated with each contrast (Cohen’s *d*_z_), we also report Bayes Factors to illustrate how statistical inferences would differ as a function of divisive vs. subtractive correction technique.

